# Poly(2-oxazoline) nanoparticle delivery enhances the therapeutic potential of vismodegib for medulloblastoma by improving CNS pharmacokinetics and reducing systemic toxicity

**DOI:** 10.1101/2020.04.30.068726

**Authors:** Duhyeong Hwang, Taylor Dismuke, Andrey Tikunov, Elias P. Rosen, John R. Kagel, Jacob D. Ramsey, Chaemin Lim, William Zamboni, Alexander V. Kabanov, Timothy R. Gershon, Marina Sokolsky-Papkov

**Author notes:** To whom correspondence should be addressed (T.R.G.); (M.SP).

## Abstract

We report a novel, nanoparticle formulation of the SHH pathway inhibitor vismodegib that improves efficacy for medulloblastoma treatment while reducing toxicity. Systemic therapies for brain tumors are complicated by restricted blood-brain barrier (BBB) permeability and dose-limiting extraneural toxicity, therefore improved delivery approached are needed. Here we show how a nanoparticle delivery system addresses these obstacles, bringing new efficacy to previously ineffective therapy. Vismodegib has been a promising agent for patients with SHH-subgroup medulloblastoma and is FDA-approved for basal cell carcinoma. However, vismodegib has limited benefit for patients with SHH-driven medulloblastoma, due to off-target toxicities and the development of resistance during therapy. We encapsulated vismodegib in polyoxazoline block copolymer micelles (POx-vismo). We then evaluated POx-vismo using transgenic mice engineered to develop endogenous medulloblastomas, testing the novel agent in a preclinical model with native vasculature and tumor microenvironment. POx-vismo showed improved CNS pharmacokinetics and reduced systemic and bone toxicity. Mechanistic studies show that POx nanoparticles did not enter the CNS, but rather acted within the vascular compartment to improve drug delivery by decreasing drug binding to serum proteins and reducing the volume of distribution. POx-vismo demonstrated improved efficacy, extending the survival of medulloblastoma-bearing mice. Our results show the potential for a simple, non-targeted nanoparticle formulation to improve systemic brain tumor therapy, and specifically to enhance vismodegib therapy for SHH-driven cancers.

## Introduction

New, targeted approaches are needed for medulloblastoma. Current treatment with surgery, radiation and chemotherapy allows most medulloblastoma patients to survive >5 years, but causes long-term neuro-cognitive injury. About one-third of medulloblastomas show SHH pathway hyper-activation, and for these patients, targeted inhibition of SHH signaling may improve therapy (*1*). Vismodegib, a small molecule inhibitor of SHH receptor component Smoothened (SMO), is FDA-approved for the treatment of basal cell carcinoma and is in clinical trials for other SHH-driven cancers (*2*). In medulloblastoma, vismodegib treatment produces initial responses, but these responses are consistently followed by recurrence (*3, 4*).

The physicochemical properties of vismodegib contribute to its limited efficacy for medulloblastoma. With extremely low aqueous solubility (0.1 µg/ml in at pH 7.0) and oral bioavailability of 31.8%, the drug is formulated for oral dosage (*5*). However, oral administration presents a challenge in pediatric population most at risk for medulloblastoma (*6*). Furthermore, Vismodegib has high affinity to serum proteins, including acid-glycoprotein and albumin, resulting in > 99% of the drug circulating in protein bound form (*5*). This low unbound fraction limits brain bioavailability; in pediatric medulloblastoma patients, the ratio of CSF vismodegib to the total drug in the plasma was 0.0026, while the ratio of CSF vismodegib to unbound drug in plasma was 50-fold higher (0.26–0.78) (*7*). These data highlight the need for optimizing vismodegib delivery by developing systemically administered formulation with reduced serum protein binding and increased brain delivery.

A variety of nanoparticles have been proposed for drug delivery to the brain. Several nanoparticle-based therapeutic products have been approved for clinical use in cancers outside the CNS, and more are currently under development (*8, 9*). Nano-formulation can enhance the solubility of hydrophobic agents, extend their systemic circulation, provide sustained drug release, enhance drug accumulation in target tissues and reduce off-site effects (*10, 11*). Polymeric micelles, formed by the self-assembly of amphiphilic block polymers, are a nanoparticle technology that has shown promise in clinical implementation (*12*). However, these systems have been limited by low drug loading and restricted tolerance for chemical structures to be loaded. In contrast, the poly(2-oxazoline) amphiphilic block copolymer (POx) forms micelles of 10-100nm, that can be fine-tuned to optimize drug loading, pharmacokinetics and biodistribution (*13*).

POx micelles have been shown to deliver poorly soluble drugs and drug combinations, improving efficacy of treatment in ovarian, breast and lung cancers (*14, 15*). We have previously shown that POx micelles can solubilize the investigational ATR inhibitor VE-821, allowing delivery into the CNS *in vivo* (*16*). To test the ability of POx delivery to optimize an FDA-approved agent with known safety profile but sub-optimal performance in prior brain tumor trials, we encapsulated vismodegib in POx micelles to generate POx-vismo. We tested POx-vismo in *G-Smo* mice which develop endogenous medulloblastoma. This transgenic model, unlike patient-derived or cell line-derived xenografts, forms tumors with native vasculature and intact blood brain barrier. Moreover, rodent xenograft models are frequently curable and have the potential to lead to overestimation of efficacy in preclinical studies. The *G-Smo* model, in contrast is highly aggressive and has not been shown to be curable with any pharmacological treatment. 100% of *G-Smo* mice develop medulloblastoma, detectable by change in head shape by P12, and, if untreated, die of tumor progression by P20. We treated medulloblastoma-bearing *G-Smo* mice with either POx-vismo or conventional vismodegib either as oral suspension or parenteral formulation, and compared pharmacokinetics, toxicity, pharmacodynamics, and efficacy.

## Results

### Self-forming POx micelles encapsulate vismodegib in nanoparticles

We prepared POx-vismo polymeric micelles using POx polymer P(MeOx_39_-b-BuOx_25_-b-MeOx_39_) (M_n_ = 8.2 kg/mol, Ð (M_w_/M_n_) = 1.11). We generated micelles with vismodegib:POx polymer ratios of 2:10, 4:10, 6:10 and 8:10 w/w (drug to polymer) using the thin film method as previously described (*14*). At all tested ratios, the loading efficiency (LE, %) of vismodegib was nearly 90% and the loading capacity was 13.5-42.4% w/w depending on the drug:polymer ratio (Table 1). POx-vismo micelles had an intensity-mean z-averaged particle size (Z_ave_) of 25-40 nm, narrow size distribution (PDI < 0.1) as determined by DLS (Fig. 1A). The POx-vismo micelles had a neutral surface charge of 8 mV (Fig. 1B). The particle size and spherical morphology of POx-vismo micelles was further confirmed by transmission electron microscopy (TEM) (Fig. 1C). The POx-vismo micelles were stable during storage at 4 °C as confirmed by HPLC and size analysis (DLS) (Fig. 1D). The POx-vismo micelles were also stable after lyophilization; lyophilized POx-vismo micelles could be re-dispersed in water, keeping their drug loading and particles size unchanged (Fig. 1E). Vismodegib was continuously released from the micelles *in vitro* under the perfect sink conditions with approximately the 25% of the drug released in 2 hours and 90% released in 12 hours (Fig. 1F).

**Table 1.** Actual vismodegib concentration, LE (%), LC (%), nanoparticles size and size distribution of POx-vismo micelles prepared at indicated drug:polymer ratios. (n = 3 ± SD)

| Ratio of drug:polymer by weight | Theoretical drug concentration (mg/ml) | Actual drug concentration (mg/ml) | LE (%) | LC (%) | z-ave diameter (nm) | PDI |
| --- | --- | --- | --- | --- | --- | --- |
| 2:10 | 2.00 | 1.57 ± 0.01 | 78.6 ± 0.5 | 13.6 ± 0.1 | 26.6 ± 2.4 | 0.07 ± 0.01 |
| 4:10 | 4.00 | 3.26 ± 0.06 | 81.6 ± 1.5 | 24.6 ± 0.5 | 34.5 ± 5.3 | 0.03 ± 0.01 |
| 6:10 | 6.00 | 4.88 ± 0.10 | 81.3 ± 1.7 | 32.6 ± 0.7 | 41.5 ± 5.9 | 0.03 ± 0.01 |
| 8:10 | 8.00 | 7.36 ± 0.15 | 92.0 ± 1.9 | 42.4 ± 0.9 | 38.3 ± 0.3 | 0.08 ± 0.01 |

**Figure 1.**
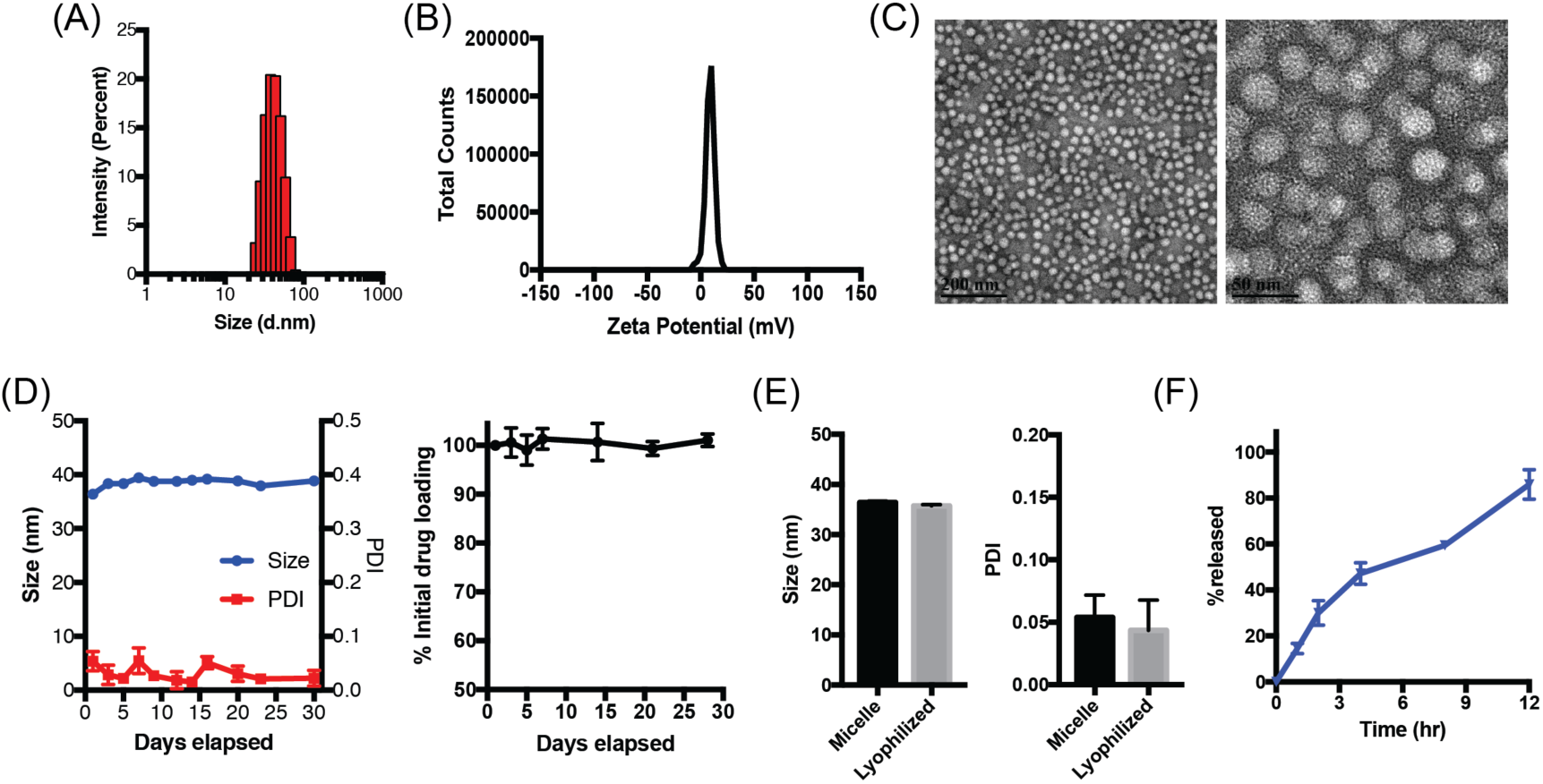
POx-vismo micelles form stable, nanometer-scale spheres. **(A)** Particle size distribution measured by DLS (z-average, Dz) **(B)** Zeta potential (*17*) and **(C)** morphology, shown by TEM. Scale bar = 200 nm (left), 50 nm (right). **(D)** Stability of the POx-vismo micelles at 4 °C as determined by actual drug measurements (left) and size distribution (right) over time. **(E)** Size of particles after reconstitution of lyophilized POx-vismo. **(F)** Vismodegib release from POx-vismo incubated in 10% fetal bovine serum (FBS) solution at 37 °C over time.

### POx micelles improve the brain and tumor delivery of vismodegib without penetrating CNS

We evaluated the effect of nanoparticle delivery on the biodistribution of vismodegib in medulloblastoma-bearing *G-Smo* mice. We administered vismodegib to *G-Smo* mice (Fig. S1), formulated as POx-vismo injected intraperitoneally (IP) or conventional vismodegib administered by either IP injection or oral gavage. We then euthanized mice after 4 hours and harvested blood, tumor and forebrain samples. We then prepared extracts of these samples and analyzed them using NMR spectroscopy methods that we developed to measure vismodegib and POx concentrations simultaneously. We identified signature peaks for vismodegib and POx that could be measured in serum and brain samples (Fig. 2A). In serum samples, vismodegib and Pox peaks increased in proportion to dose (Fig. 2A insets)

**Figure 2.**
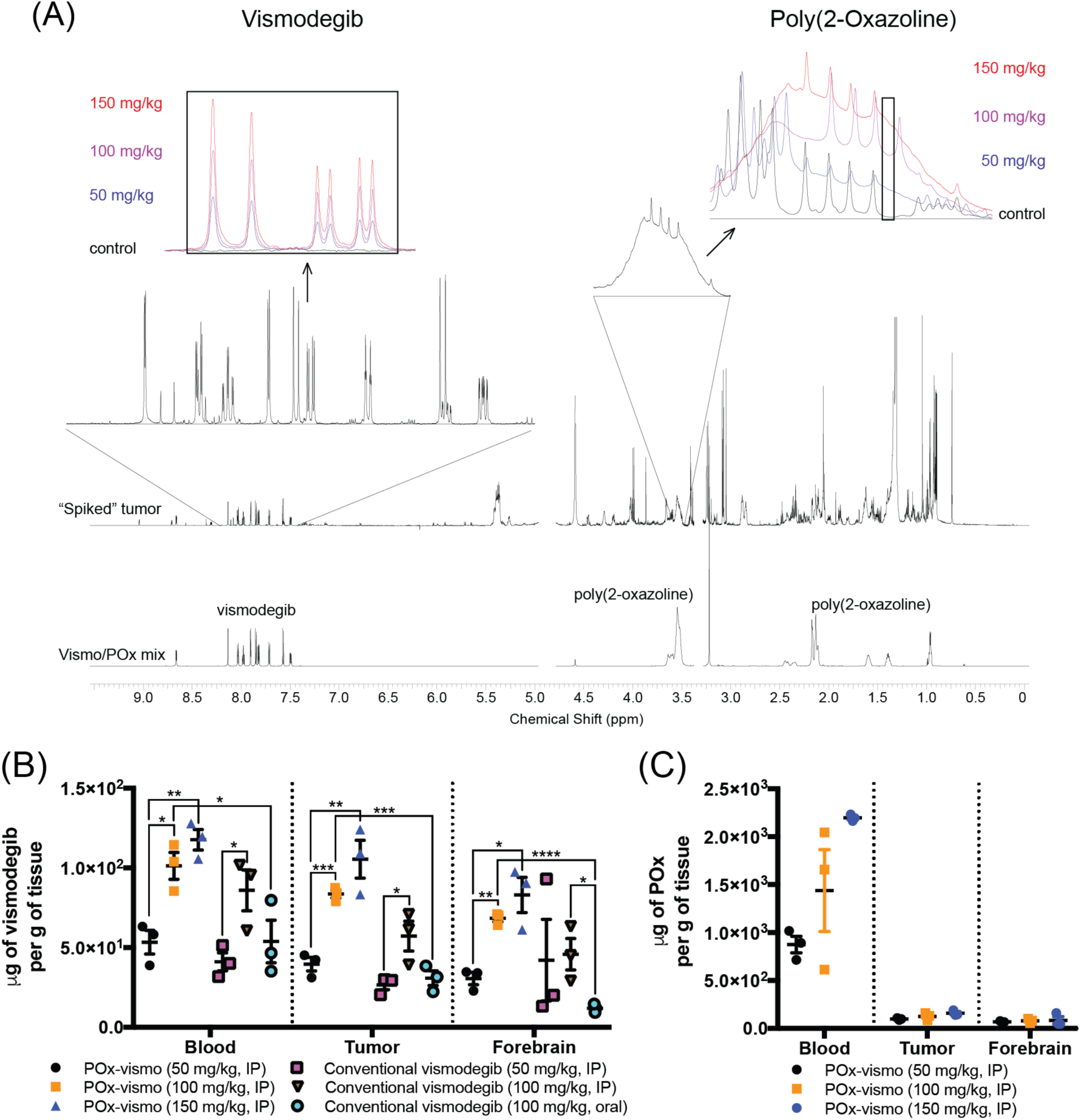
Differential distribution of vismodegib and POx components of POx-vismo in the vascular and CNS compartments. **(A)** NMR spectra with the signature peaks for vismodegib and POx highlighted. Insets show dose-dependent changes in regions that correspond to vismodegib or POx in representative spectra from serum samples. Note that POx peak is a broad peak that spans several, more narrow peaks for unrelated molecules. To maximize specificity, we integrated an area within the broad POx peak that falls between unrelated peaks. The area under this portion of the curve was proportional to POx concentration over the range of tested standards. **(B)** Vismodegib concentrations in serum, tumor and forebrain after administration of the indicated formulation and dose **(C)** POx concentration in serum, tumor and forebrain following IP injection of the indicated doses of POx-vismo. *p < 0.05, **p < 0.005, ***p < 0.0005. For B and C, dots indicate data from individual replicates and error bars are the SEM.

We compared vismodegib in blood, tumor and forebrain samples from mice treated with each formulation (Fig. 2B). Escalating POx-vismo doses of 50 mg/kg, 100 mg/kg and 150 mg/kg produced a linear increase in vismodegib concentration in blood, tumor and forebrain, and IP conventional vismodegib showed a similar pattern. Systemic administration of conventional vismodegib resulted to comparable levels to POx-vismo in plasma at the same dose of 100 mg/kg. Oral vismodegib produced significantly lower vismodegib concentrations in blood, tumor and forebrain compared to POx-vismo and showed a similar trend toward lower concentrations compared to systemic vismodegib that was statistically significant in the forebrain compartment. These data show that oral administration produced similar or inferior concentrations compared to parenteral administration.

NMR spectroscopy showed that the POx polymer component of POx-vismo, unlike the vismodegib component, did not penetrate the BBB. While escalating doses of POx-vismo produced increasing concentrations of POx in the blood, POx concentrations in the tumor and forebrain remained below the threshold of detection at all tested dose (Fig. 2C). Based on these data, we conclude that the nanoparticle carrier dissociates from the vismodegib payload prior to the entry of vismodegib into the CNS.

To determine the effect of POx-vismo delivery on pharmacokinetics, we compared vismodegib concentrations in the blood, medulloblastoma and forebrain of tumor-bearing *G-Smo* mice at different intervals after parenteral administration of either POx-vismo or conventional vismodegib. We injected *G-Smo* mice on P10 with a single IP injection of either POx-vismo or conventional vismodegib, both dosed at 100 mg/kg vismodegib. We then harvested serum, tumor and brain tissues at 5 minutes, 2 hours, 4 hours, 8 hours or 24 hours after administration and measured vismodegib concentrations by Liquid chromatography–mass spectrometry (LC-MS). In sagittal brain sections from replicate mice harvested 4 or 24 hours after administration, we used infrared matrix assisted laser desorption electrospray ionization (IR-MALDESI) imaging to determine the spatial distribution of vismodegib across tissue slices containing both medulloblastoma and brain.

LC-MS analysis showed that POx-vismo produced higher, more long-lasting vismodegib accumulation in all tissues, with a higher overall tissue drug exposure. POx-vismo induced significantly higher vismodegib concentrations in serum, medulloblastoma and forebrain at 2, 4, and 8 hours, compared to conventional vismodegib (Fig. 3A-C). At 24 hours, vismodegib was below the level of detection in both tumor or brain, preventing comparison. Consistent with higher peaks and more sustained concentrations, the total tumor vismodegib exposure, measured as area under the curve (AUC) was significantly higher in tumor, adjacent brain and serum) following administration of POx-vismo compared to conventional vismodegib (Fig. 3D). POx- vismo-treated mice also showed higher vismodegib concentration ratios for tumor:serum and forebrain:serum (Fig. 2E, F), indicating improved distribution across BBB. Along with increased tumor drug exposure, nanoparticle formulation decreased the calculated vismodegib volume of distribution (Vd) (Fig. 3G) and clearance (CL) (Fig. 3H), indicating reduced distribution to non-target organs and improved the retention of the drug in the target tissues. Analysis of vismodegib distribution in non-tumor mice, showed similar results (Fig. S2), indicating that the improved CNS penetration of vismodegib administered as POx-vismo, and reduced systemic distribution, were not due to tumor-specific changes in the BBB.

**Figure 3.**
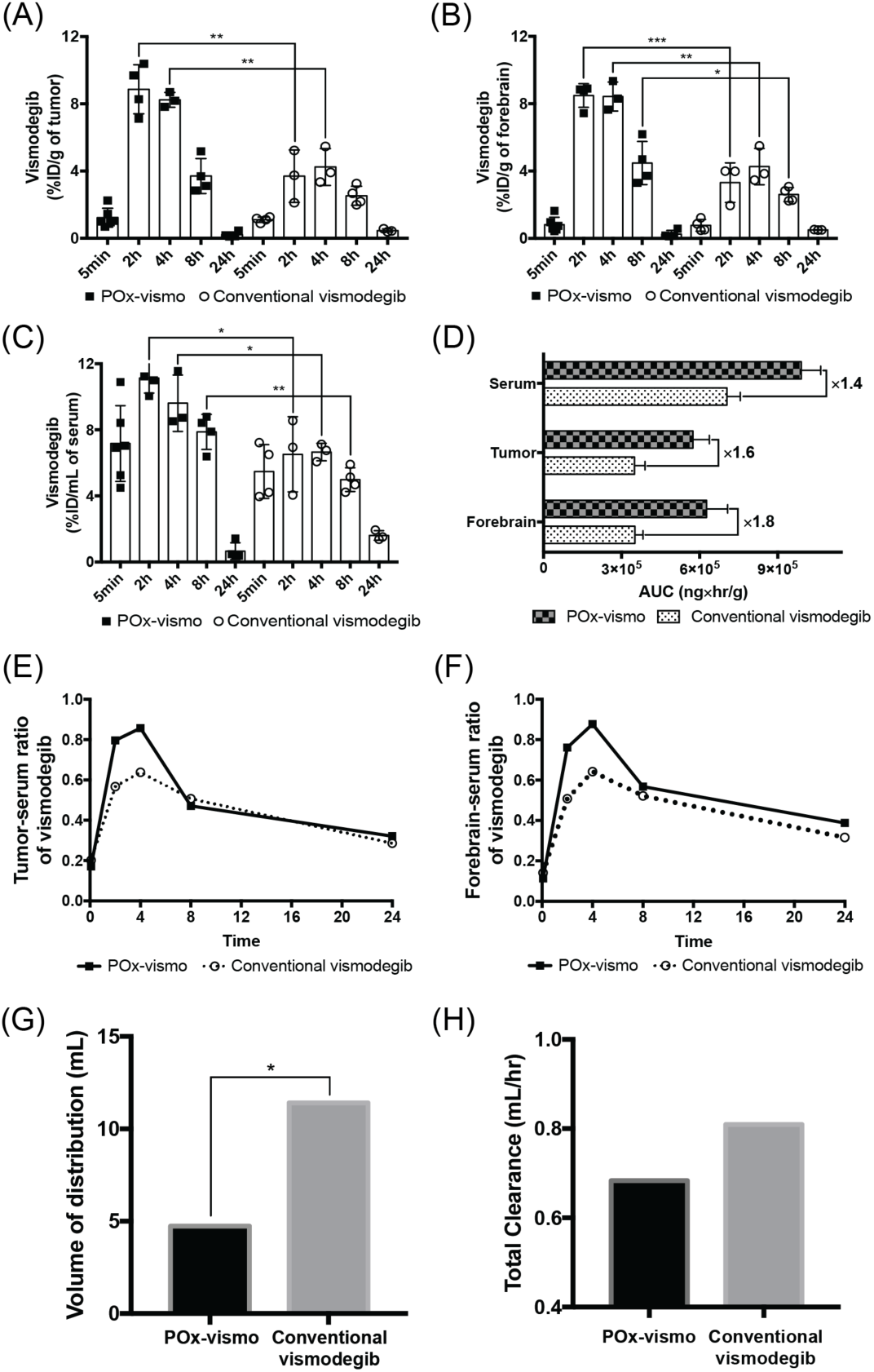
Pharmacokinetic profile of POx-vismo and conventional vismodegib in medulloblastoma-bearing *G-Smo* mice. Dot plots of vismodegib concentration in (A) tumor, (B) forebrain and (C) serum. (D) AUC comparison of serum, tumor, and forebrain, (E) Tumor-serum ratio of vismodegib, (F) forebrain-serum ratio of vismodegib, (G) volume of distribution and (H) total clearance of vismodegib following administration of 100 mg/kg of POx-vismo or conventional vismodegib in tumor bearing mice. *p < 0.05, **p < 0.005, ***p < 0.0005. For A-C, data are expressed as dot plots with column (means ± SEM), the number of each group is expressed as the number of dots (n ≥ 3).

We analyzed the spatial distribution of vismodegib across sections of tumor and brain using IR-MALDESI (*18*). This technique uses infrared laser scanning across frozen tissue sections to excite the overlying matrix of ice, causing desorption of endogenous and exogenous ions that can be detected mass spectrometry. IR-MALDESI quantifies the concentrations of species at all detected molecular weights with spatial resolution, generating ion-specific heatmaps. We validated this technique by imaging endogenous cholesterol, which we found to be at relatively high concentration in normal brain compared to tumor (Fig. 4A, B). MALDESI detected vismodegib evenly distributed in tumor and forebrain 4 hours after IP injection of either POx-vismo or conventional vismodegib (p=0.630; Fig. 4B, C). Importantly, IR-MALDESI, in contrast to LC-MS, was able to measure vismodegib concentrations at 24 hours; we attribute this increased sensitivity to the lack of an extraction step. POx-vismo treated mice showed higher vismodegib signal at both 4h and 24h post injection (Fig. 4B, C).

**Figure 4.**
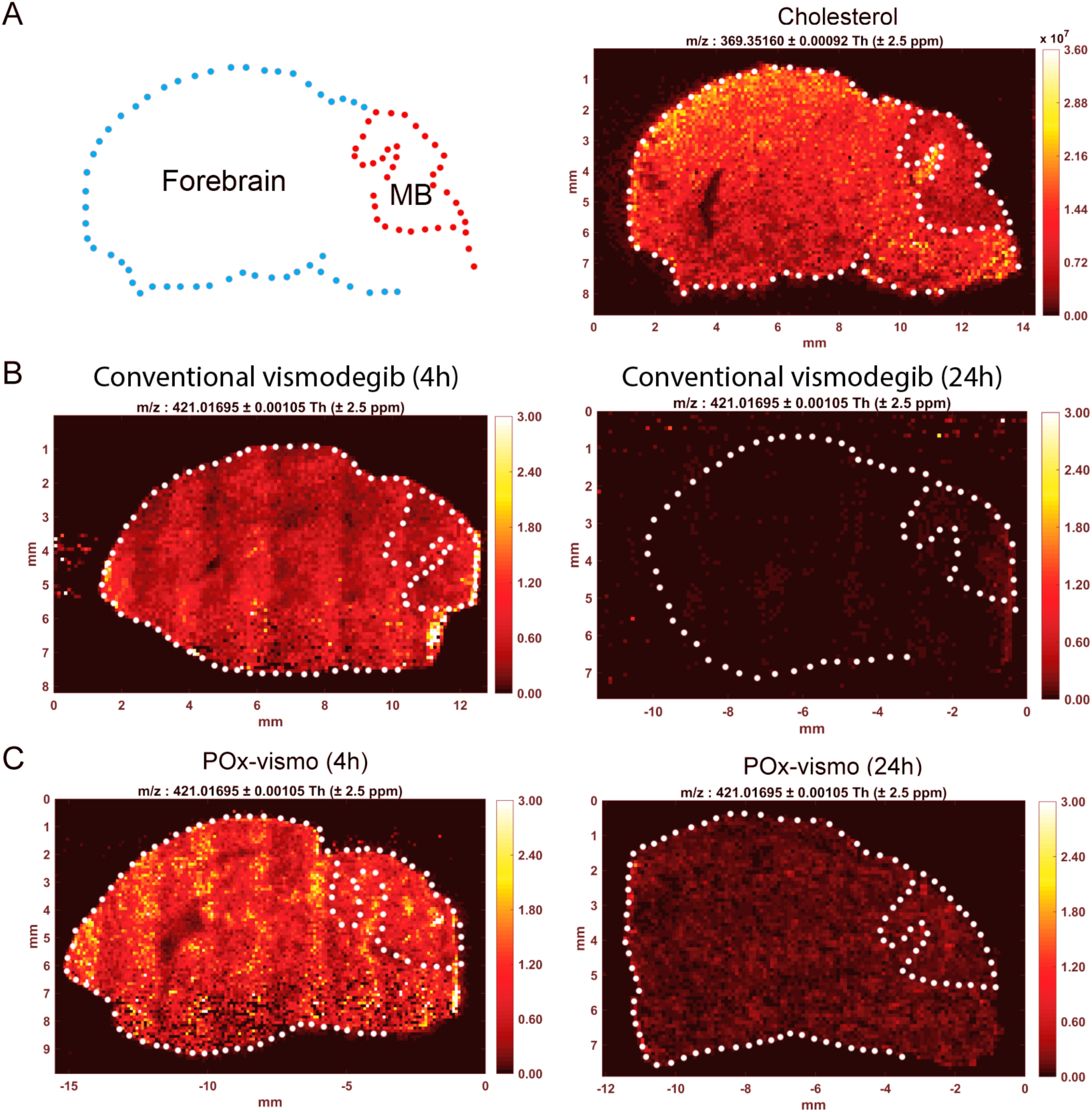
Widespread vismodegib distribution in the brain, with increased retention after POx-vismo administration, demonstrated by IR-MALDESI. **(A)** Left panel, contour of brain sample showing forebrain and medulloblastoma (MB) and right panel, IR-MALDESI analysis of cholesterol in a control brain (m/z :369). **(B)** IR-MALDESI analysis of vismodegib (m/z: 421) in representative sagittal brain sections from *G-Smo* mice following a single dose or conventional vismodegib at 4 hours (left) and 24 hours (right) **(C)** IR-MALDESI MSI analysis of vismodegib (m/z: 421) as in **(B)** following a single dose of POx-vismo at 4 hours (left) and 24 hours (right).

### Nanoparticle carrier reduces vismodegib binding to serum albumin and improves bioavailability

The exclusion of the nanoparticle carrier from the CNS suggested that the mechanism of nanoparticle effect may occur in the blood. As the penetration of vismodegib into the brain is known to be affected by its binding to serum protein, we determined if POx formulation changes the serum protein binding of vismodegib. We compared albumin binding of vismodegib in POx- vismo and conventional formulations by measuring the quenching of the fluorescence of albumin tryptophan (*19*). The fluorescence quenching was lower for samples incubated with POx-vismo versus conventional vismodegib, indicating that POx formulation reduced albumin binding (Fig. 5A, B). Conversely, the intensity ratio at 340 nm was consistently higher for POx-vismo at each step in the increase of drug (Fig. 5C). Next, we evaluated protein binding over time. After 30 minutes, over 30% of total vismodegib remained unbound, decreasing to 10% over 4 hours. In contrast, the unbound fraction of conventional vismodegib was marginal at 30 minutes. This is consistent with the improved tumor:serum and forebrain:serum ratio measured at early time points (up to 4h) following administration of either POx-vismo or conventional vismodegib (Fig. E,F). These data show that POx nanoparticle encapsulation reduces vismodegib protein binding, suggesting a mechanism for improved CNS penetration of vismodegib in POx-vismo.

**Figure 5.**
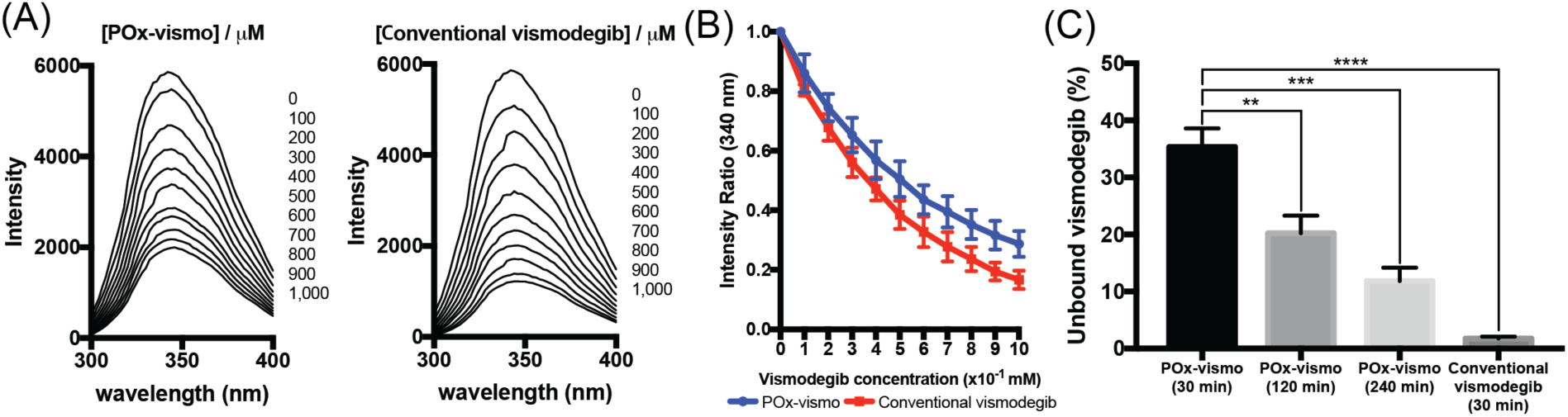
Reduced vismodegib:protein binding with POx-vismo. **(A)** Tryptophan fluorescence quenching assay by addition of either POx-vismo or conventional vismodegib at given concentration of vismodegib (right), **(B)** the fraction of unbound vismodegib by addition of either Pox-vismo or conventional vismodegib in diluted FBS solution. **(C)** Unbound vismodegib after the incubation in FBS. *p < 0.05, **p < 0.005, ***p < 0.0005. Each point is mean ± SEM.

### POx-vismo reduces systemic toxicity of vismodegib

To determine whether nanoparticle delivery altered the systemic toxicity of vismodegib, we compared healthy P10 mouse pups treated with either PO-vismo or conventional vismodegib. To evaluate systemic toxicity, we compared growth of pups on extended regimens of POx-vismo or conventional vismodegib to that of untreated age-matched littermate controls (Fig. 6A). POx-vismo was injected IP and conventional vismodegib was administered either IP or by oral gavage. Drug was administered on postnatal days 10-12 and then every other day until P35. All mice including littermate controls were weighed daily. Mice treated with 100 mg/kg POx-vismo gained weight similar to untreated controls. In contrast mice on oral or IP conventional vismodegib at 100 mg/kg showed significantly decreased weight gain that required a 50% increased dose of 150 mg/kg POx-vismo to replicate. These studies show that POx formulation reduced systemic toxicity.

**Figure 6.**
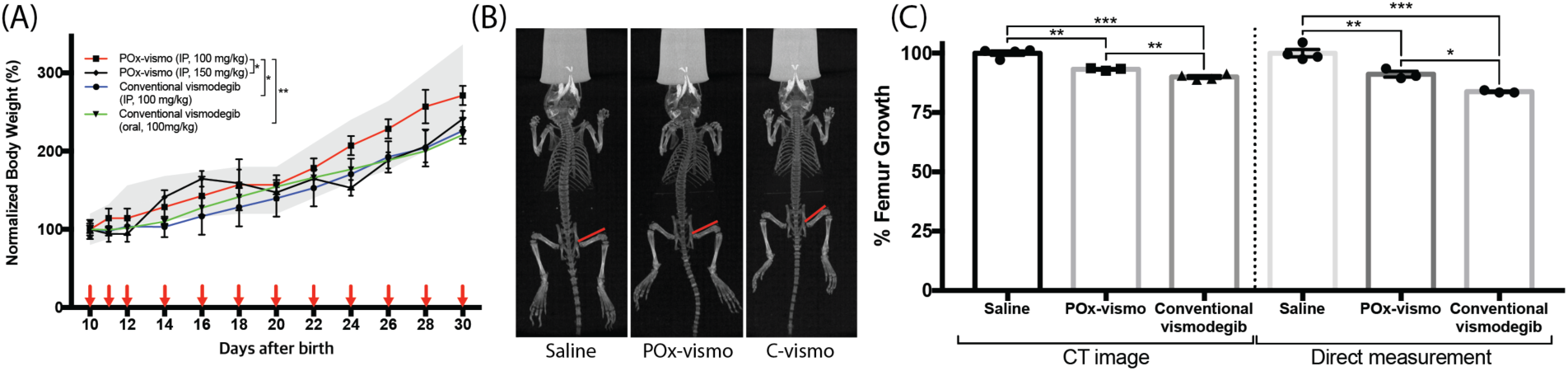
Reduced toxicity in POx-vismo treated mice. (A). The weights of mice treated with the indicated formulations are graphed over time. The gray range indicates the mean weights ± SEM of littermate controls. (B) Whole body CT scan of mice treated as indicated. Red lines show the femur length definition used for measurements. (C) femur growth, normalized to the mean for saline-treated controls, as determined by CT scan, or by direct measurement. *p < 0.05, **p < 0.005, ***p < 0.0005. Each point is mean ± SEM.

Vismodegib is known to impair bone growth, and we compared the bone toxicity in mice treated with POx-vismo or conventional vismodegib (*20, 21*). We treated P10 mice with 4 daily doses of 100 mg/kg of either formulation, administered IP, then performed CT scans and measured femur length, as in prior vismodegib toxicity studies (Fig. 6B). We also isolated and directly measured the length of the dissected femurs. These studies showed that both POx- vismo and conventional vismodegib reduced femur length, as in prior studies, but that POx- vismo caused significantly less growth reduction (Fig. 6C).

### POx-vismo and conventional vismodegib show similar pharmacodynamic effects

Next, we examined whether nanoparticle formulation altered the pharmacodynamic effect within the CNS. We treated *G-Smo* mice with a single 100 mg/kg dose of POx-vismo administered IP or conventional vismodegib, administered either IP or orally. We then harvested tumors after 24 hours and compared the fractions of cells in tumors that expressed the proliferation marker phosphorylated RB (pRB; Fig. 7A). POx-vismo and conventional vismodegib IP similarly suppressed pRB (p=0.9987), while oral vismodegib showed less suppression, approximating the effect of IP vismodegib at 50% lower dose (Fig. 7B). To compare pharmacodynamic effects of repeated doses, we examined tumor pathology after 3 daily IP doses of either POx-vismo or conventional vismodegib. We again found similar suppression of proliferation, with reduced tumor size and expression of both pRB and Proliferating Cell Nuclear Antigen (PCNA) (Fig. 7B). These data show that POx-vismo suppressed SHH-driven proliferation more effectively than the oral formulation and at least as effective as systemically administered conventional vismodegib.

**Figure 7.**
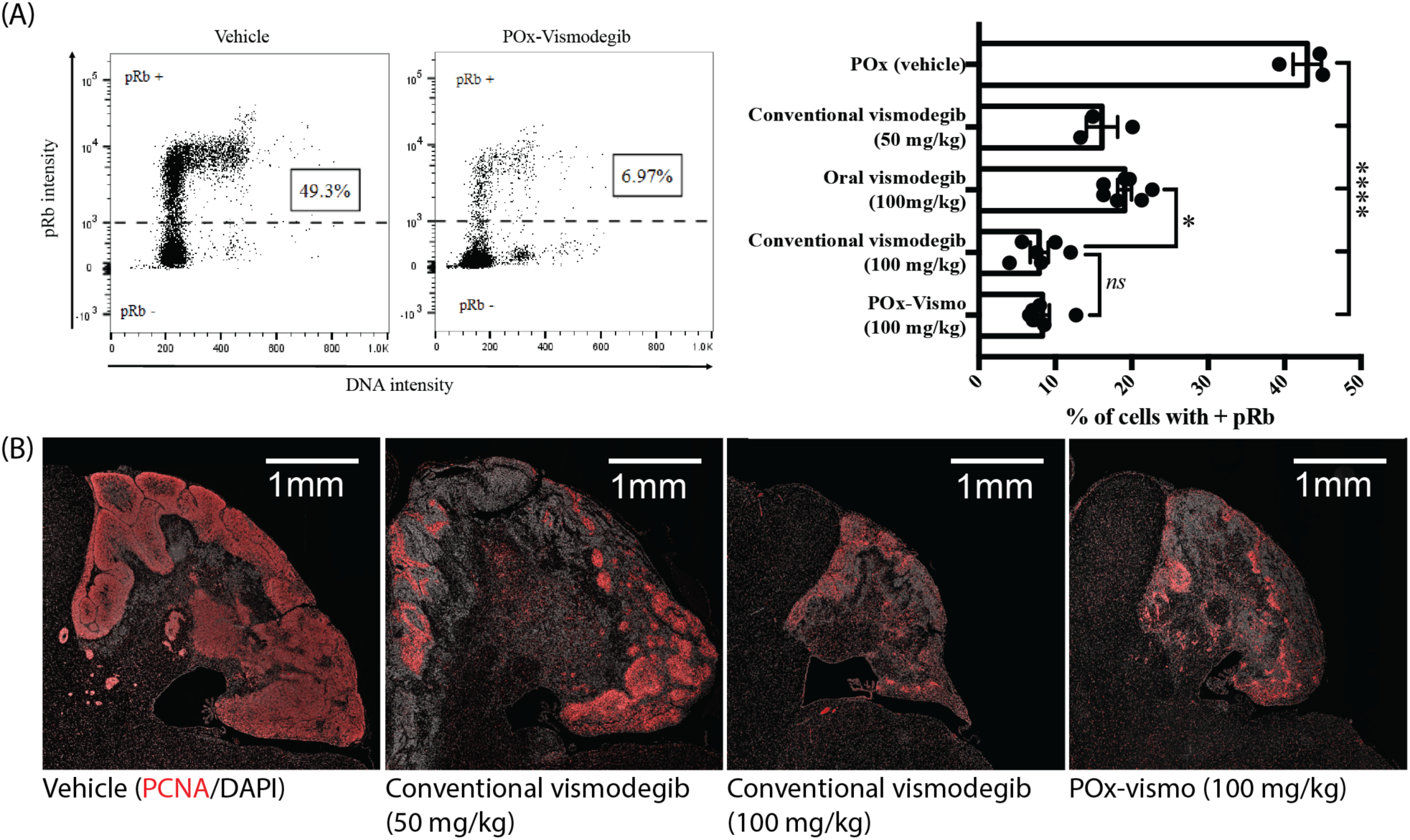
Pharmacodynamic response to POx-vismo and conventional vismodegib. **(A)** Flow cytometry quantification of pRB^+^ cells in untreated tumors and in tumors 24 hours after a single IP administration of POx-vismo, or conventional vismodegib, administered IP or orally, compared to POx vehicle-injected controls. Cells of representative replicates treated as indicated are plotted according to pRB intensity versus DNA content. The dotted line indicates the threshold of detection for pRB. The pRB^+^ fractions for all replicates in each treatment group are graphed to the right, with columns indicating the means and error bars indicating the SEM. **(B)** PCNA immunofluorescence staining of sagittal medulloblastoma sections from representative P13 *G-Smo* mice, 24 hours after three daily IP injections of the indicated formulation. Nuclei are counterstained with DAPI.

### POx-vismo micelles prolong survival in GEMM model of SHH medulloblastoma

To determine whether improved pharmacokinetics and toxicity profile of POx-vismo improved the therapeutic efficacy of vismodegib, we compared the survival of *G-Smo* mice on a regimen of either POx-vismo or conventional vismodegib administered IP or orally. Our prior studies showed conventional vismodegib initially suppressed proliferation in SmoM2-driven medulloblastomas but failed to extend mouse survival (*18*). We randomized *G-Smo* mice to 4 groups: conventional vismodegib 100 mg/kg oral, conventional vismodegib 100 mg/kg IP, POx-vismo at 100 mg/kg IP and saline-injected controls. The survival time to the humane endpoint was considered event-free survival (EFS). All *G-Smo* mice in the saline-injected control group survived less than 20 days, consistent with our prior data (*22, 23*). Treatment with treatment with 100 mg/kg POx-vismo significantly improved the EFS (p < 0.001 Log-rank test) with 30% of mice surviving to 35 days (Fig. 8A). In contrast, 100 mg/kg conventional vismodegib administered either IP or orally, failed to extend the EFS. To determine whether the failure of conventional vismodegib to extend the survival was due to systemic toxicity, we treated another group of mice with conventional vismodegib 50 mg/kg IP. Although this dose produced a detectable suppression of pRB (Fig. 7A), there was no benefit in survival time. These data show that POx vismo produced superior efficacy compared to conventional vismodegib.

**Figure 8.**
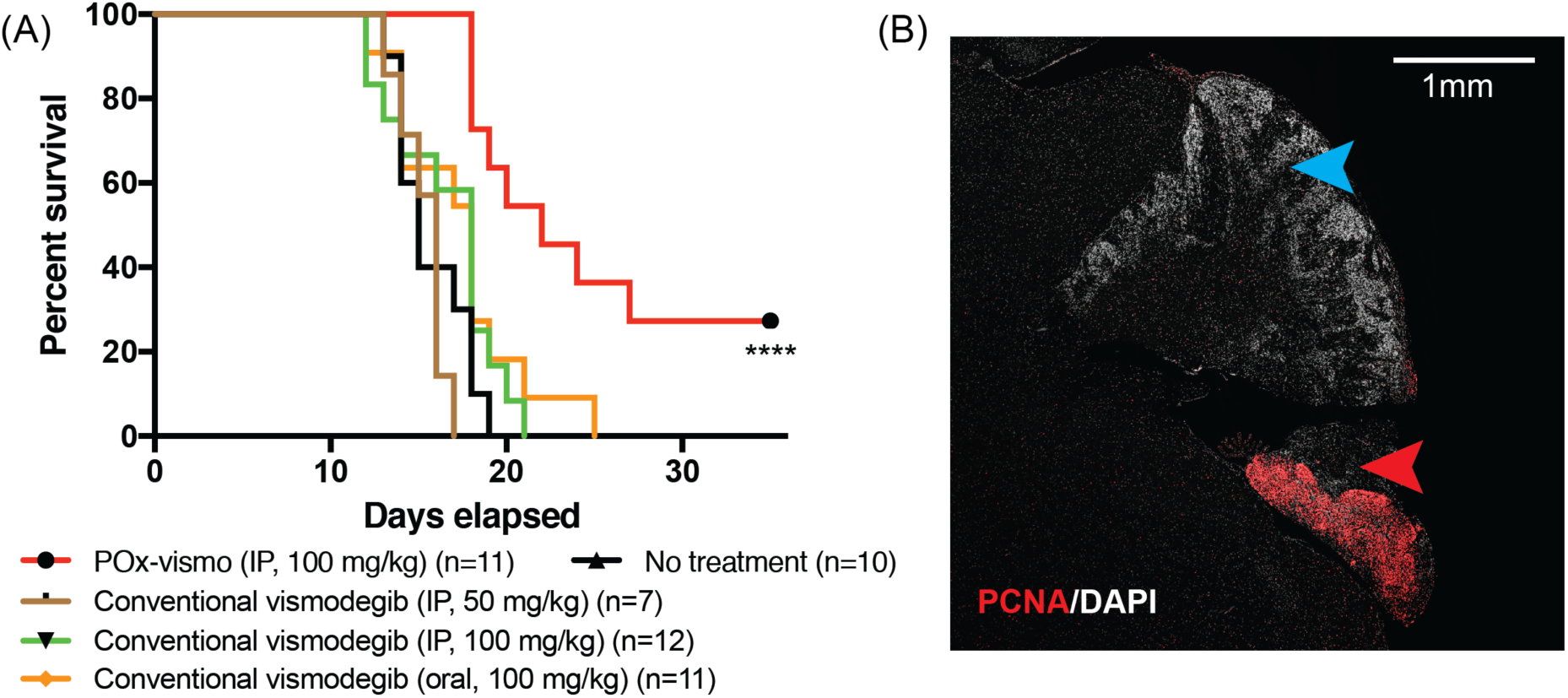
Increased efficacy of POx-vismo compared to conventional vismodegib. **(A)** Kaplan-Meier survival curves for *G-Smo* mice treated with the indicated regimen, compared to no treatment; *** indicates p < 0.001 (vs. No treatment). All other curves were not significantly different from No treatment. **(B)** PCNA immunofluorescent staining (red), in a sagittal medulloblastoma section from a representative P35 *G-Smo* mouse, treated with 100 mg/kg of POx-vismo (P35). Nuclei are counterstained with DAPI. Red arrowhead indicates a region of proliferative tumor. Blue arrowhead indicates a differentiated, non-proliferative region.

We evaluated the tumor pathology in POx-vismo-treated mice that survived beyond the 20-day maximum EFS observed in the other groups. In each of these mice, we found regions of highly proliferative cells within the cerebellum, interspersed with regions of differentiated, non-proliferative cells (Fig. 8B). The non-proliferative regions indicate that vismodegib effectively interrupted SHH-driven proliferation and restricted tumor growth, while the proliferative regions show that even in the longer-term survivors, residual disease persisted. While POx-vismo was not curative, no known drug is curative as a single agent for medulloblastoma. POx-vismo was singularly effective in extending the life of *G-Smo* mice, while conventional vismodegib, oral or parenteral, failed to provide a survival benefit.

## Discussion

Vismodegib has shown clinical efficacy for treatment of SHH-driven cancer outside the CNS, but has not been as effective for SHH subtype medulloblastoma either in animal models (*18*) or in patients (*3, 24*). We now show that optimizing vismodegib delivery using nanoparticles markedly reduces toxicity and improves efficacy, providing strong support for the use of the POx nanoparticle platform. POx-vismo formed uniform nanometer-scale particles with high loading capacity that were stable in aqueous media and could by lyophilized and reconstituted with water, facilitating clinical implementation.

Our analyses show that POx formulation increased drug exposure in the CNS, in both medulloblastoma and forebrain and reduced systemic exposure, providing an explanation for both improved anti-tumor effect and reduced toxicity. Although low doses of vismodegib can induce a pharmacodynamic response, exposure to sub-optimal concentrations of cytotoxic drugs is associated with development of resistance and failure to respond to treatment (*25, 26*). Resistance to vismodegib develops rapidly during therapy and was identified as main reason for treatment failure (*27, 28*). We were able to achieving higher, more sustained drug concentration in the target tumor tissues, with lower toxicity using POx-vismo, producing improved antitumor efficacy compared to conventional vismodegib.

Our NMR studies provide important insight into the mechanism through which POx nanoparticle encapsulation improves pharmacokinetics. NMR simultaneously detected the polymer carrier and the drug, without the need to chemically modify their structures, which can affect biodistribution (*29*). We found that the vismodegib component of POx-vismo enters the brain and brain tumor, while the POx component does not. This finding indicates that the nanoparticle carrier releases its payload outside the BBB and underscores the importance of the effect of the nanoparticle carrier on drug dynamics in the blood compartment. Nanoparticle encapsulation reduced vismodegib serum protein binding, and free vismodegib is known to pass through the BBB. Taken together, our data demonstrate a mechanism in which POx nanoparticle delivery improves drug penetration into normal brain and brain tumors by increasing the fraction of drug available for passage across the BBB. Importantly, since the nanoparticle does not enter into the CNS, there is reduced potential that it will cause untoward effects on the normal brain.

Our work advances the development of nanoparticle drug delivery for brain tumors by working entirely in a primary, genetic model in which tumors grow in the intracranial space, with an endogenous BBB. Diverse nanoparticles have shown efficacy against medulloblastoma cell lines either *in vitro* or in xenografts (*30*), including poly(lactic-co-glycolic acid) conjugated to polyethylene glycol (PLGA-PEG)-based nanoparticles delivering the SHH pathway inhibitor HPI-1 (*31*). One nanoparticle system tested in an endogenous preclinical brain tumor model, iron oxide nanoparticles-coated with chitosan and polyethyleneimine-PEG copolymer, coated with the tumor-targeting chlorotoxin peptide, effectively delivered siRNA in a primary mouse glioma model (*32, 33*). Our study opens new potential for brain tumor nanoparticle-drug delivery by demonstrating that a drug that is FDA-approved and in clinical use can be made more effective and less toxic. Importantly, POx micelles are a versatile system that can efficiently load diverse small molecules. Moreover, the non-targeted nature of POx micelles may be advantageous by not requiring a specific receptor that cancer cells can down-regulate to become resistant.

Our data clearly show that in mice POx-vismo is less a toxic and more effective, parenteral alternative to conventional vismodegib. Phase 1 trials to test the safety of POx-micelles loaded with other agents are currently on-going and early results of one phase 1 trial showed that poly(2- oxazoline) nanoparticles similar to POx micelles in this study were well tolerated (*34*). The improved efficacy of POx-vismo and the need for more effective treatments for SHH-driven cancers support the testing of POx-vismo in humans.

The persistence of small rests of proliferating medulloblastoma cells in mice treated with POx-vismo through P35 highlights the need for combinations of pharmacologic agents to forestall resistance. Vismodegib-resistant cells may have different, specific sensitivities that may be defined in future studies of up-regulated signaling pathways in recurrent tumors (*35*). The versatility of the POx system as a potential carrier for diverse agents allows for delivery of vismodegib combined with other specific inhibitors or chemotherapeutic agents. Our data show that optimization through nanoparticle formulation can improve the therapeutic index and efficacy of systemically administered pharmaceutical agents for brain tumor therapy, with the potential to improve survival and quality-of-life for brain tumor patients.

## Materials and Methods

### Materials

All materials for the synthesis of poly(2-oxazoline) block copolymer, N-Methyl-2- pyrrolidone (NMP), PEG300, FBS and acetone were purchased from Sigma Aldrich (St. Louis, MO). Vismodegib free base was purchased from LC Laboratories (Woburn, MA). For the parenteral administration of conventional vismodegib, vismodegib was dissolved in NMP and then diluted in PEG300 to a final vismodegib concentration of 21 mg/mL.

Water and acetonitrile (HPLC grade) were purchased from Fisher Scientific Inc. (Fairlawn, NJ). PCNA antibody for immunohistochemistry was purchased from Abcam (Cat# ab92552; Cambridge, MA). Alexa-647-conjgated antibodies to pRB were purchased from Cell Signaling Technology (cat# 8974; Danvers, MA). To prepare tumors for flow cytometry, papain was purchased from Worthington Biochemical Corporation (Lakewood, NJ) and Fix&Perm® cell fixation and permeabilization kit was purchased from Thermo Fisher Scientific.

### Preparation of POx-vismo micelles

The amphiphilic triblock copolymer (P(MeOx_39_-b-PBuOx_25_-b-PMeOx_39_)), Mn = 8.2 kg/mol, PDI = 1.11)) was synthesized and characterized as previously described (*14*). Vismodegib-loaded polymeric micelle formulation (POx-vismo) was prepared by the thin film hydration method (*36*). Briefly, stock solutions of the polymer and vismodegib (10 mg/ml in acetone) were mixed together at the pre-determined ratios (2:10-8:10 drug to polymer w/w ratios). The organic solvent was evaporated at 60 °C under a stream of nitrogen gas to form a thin-film of drug-polymer homogenous mixture. To obtain well-dried thin film, the films were dried in the vacuum chamber (approx. 0.2 mbar) overnight. Next, the thin films were rehydrated with saline and then incubated at 60 °C for 10 min to self-assembly into drug-loaded polymeric micelles solution. The formed POx-vismo micelles were centrifuged at 10,000 rpm for 3 minutes (Sorvall Legend Micro 21R Centrifuge, Thermo Scientific) to remove non-loaded vismodegib.

### Characterization of POx-vismo micelles

The Z_ave_ and the polydispersity index (PDI) of POx-vismo were determined using a Nano-ZS (Malvern Instruments Inc., UK) dynamic light scattering (DLS). Each sample was diluted with saline to yield 1 mg/mL final polymer concentration. Z_ave_ and the polydispersity index (PDI) of POx-vismo were determined by cumulate analysis. Results are the average of three independent micelle samples measurements.

The morphology of POx-vismo micelles was determined using a LEO EM910 TEM operating at 80 kV (Carl Zeiss SMT Inc., Peabody, MA). The micelles were stained with negative staining (1% uranyl acetate) before imaging. Digital images were obtained using a Gatan Orius SC1000 CCD Digital Camera in combination with Digital Micrograph 3.11.0 software (Gatan Inc., Pleasanton, CA).

The final concentration of vismodegib in POx-vismo micelles was determined by HPLC (Agilent Technologies 1200 series) using Agilent eclipse plus C18 3.5 μm column (4.6mm × 150mm) with a mixture of acetonitrile/water (30%/70% v/v, 0.01% trifluoroacetic acid mobile phase. The samples were diluted with mobile phase to final concentration of 100 μg/mL of vismodegib (and injected (10 μL) into the HPLC system. The flow rate was 1.0 mL/min, and column temperature was 40 °C. Detection wavelength was 245 nm. Vismodegib concentration was quantified against free vismodegib analytical standards.

Loading efficiency (LE) and loading capacity (LC) calculations. The following equations were used to calculate LE and LC of vismodegib in POx-vismo micelles:

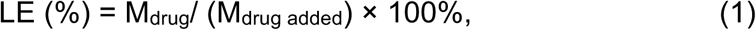

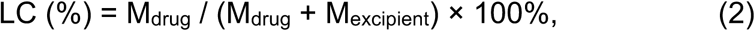

Where M_drug_ and M_excipient_ are the mass of the solubilized drug and polymer excipient in the solution, while M_drug added_ is the weight amount of the drug added to the dispersion during the preparation of the micelle formulation.

### Nanoparticle stability

The POx-vismo micelles solutions (10 mg/mL of polymer concentration) were incubated in saline at 4 ^°^C for 1 month. At pre-determined time points the aliquots were removed and the vismo concentration was measured by HPLC and the Z_ave_ and PDI of the POx-vismo were measured by DLS as previously described.

### Stability to lyophilization

Freshly prepared POx-vismo micelles solution (100 μL, 1 mg/mL of vismodegib in saline) was immediately frozen in liquid nitrogen for 5 min and lyophilized to obtain white powder of POx-vismo. Powder formulation of POx-vismo was resuspended in DI water to form POx-vismo micelles solution, centrifuged (10,000 rpm and 3 minutes) to remove any drug or polymer precipitate. Vismo concentration was measured by HPLC and the Z_ave_ and PDI of the POx-vismo were measured by DLS as previously described.

### Release of vismodegib from POx-vismo micelles

Release of vismodegib from POx-vismo micelles was investigated using membrane dialysis method against 10% solution of BSA in phosphate buffered saline (pH 7.4) at 37 C. Briefly, POx-vismo was diluted in saline to final concentration of 0.6 mg/mL of vismodegib. Subsequently, 100 μL of the diluted POx-vismo solutions were loaded into floatable Slide-A-Lyzer MINI dialysis devices (500 µL capacity, 3.5 kDa MWCO; Thermo Fisher Scientific). The dialysis devices (n=3) were floated in 20 mL of 10% BSA in PBS solution in compliance with the perfect sink conditions requirements. At each time point the samples were withdrawn from dialysis devices and the amount of vismodegib was quantified by HPLC. Drug release profiles were constructed by plotting the percentage of vismodegib released from POx-vismo micelles over time.

### Mouse breeding

Medulloblastoma-prone *G-Smo* mice were generated from the cross between *hGFAP-Cre* (generously shared by Dr. Eva Anton, UNC) and *SmoM2*^*loxP/loxP*^ (Jackson labs, Stock #005130) mouse lines. All mice were of species *Mus musculus* and crossed into the C57BL/6 background through at least five generations. All animal studies were carried out with the approval of the University of North Carolina Institutional Animal Care and Use Committee under protocol 16-099.

### NMR analysis

*G-Smo* mice were injected with 50-150 mg/kg POx-vismo intraperitoneally (IP) and conventional vismodegib was administered by either IP injection (50-100 mg/kg) or oral gavage (100mg/kg) on P12. 4h post injection, 3 replicate mice were euthanized. Blood and brains (forebrain and tumor tissue), were collected. 600 µL of 100% acetonitrile was added to the frozen tissue, then tissue was homogenized with pestle, frozen and thawed twice. Next, 400 µL of water was added, sample was vortexed, frozen and thawed again. Supernatant was separated by centrifugation and lyophilized. The powder was resuspended in 600 µL deuterated methanol. The ^1^H NMR spectra were acquired at 25 °C using a 19.97 T Brucker spectrometer equipped with a 5 mm HCN NMR probe with a one-pulse sequence using a 90° flip angle, with a 1.5 s presaturation pulse on residual water, a 2.5 s acquisition time and 1.5 s relaxation time resulting in a 4 s repetition time. The sweep width was 6,000 Hz and acquired with 15,000 complex points, and 128 transients. All NMR spectra were processed using ACD/1D NMR Manager software (version 12.0; Advanced Chemistry Development). Imported FIDs were zero-filled to 32,000 points, and an exponential line broadening of 0.3 Hz was applied before Fourier transformation. Phase and baseline correction were conducted for the spectra. Peaks were integrated and values were exported to Excel for data processing.

### Pharmacokinetic analysis

4 replicate *G-Smo* mice were used for each dose and formulation. The mice were injected IP with 100 mg/kg of either POx-vismo or conventional vismodegib on P12. At indicated sampling times, mice were euthanized. Blood and brains (forebrain and tumor tissue) were collected. Samples were spiked with internal standard (reserpine) solution and calibration standard solution, treated with 1% formic acid in acetonitrile and then centrifuged to obtain the supernatant. The acetonitrile was evaporated and the reconstitution solution (acetonitrile/water/0.1% formic acid) was added to each sample. The samples were centrifuged again and the supernatant was analyzed by LC-MS. Separation was carried out on a HALO PFP column at 40 °C (2.1×50 mm, 5 µm; Advanced Materials Technology, Wilimington, DE, USA) using a Shimadzu UFLC system (PAL system, Lake Elmo, MN, USA). Samples were eluted with water and acetonitrile (containing 0.1% of formic acid) (90:10, v/v) at a flow rate of 0.5 mL/min. The MS was performed on Thermo TSQ Quantum Ultra (Thermo Fisher Scientific, Waltham, MA, USA) with an atmospheric pressure chemical ionization (APCI) source in positive ion mode operated with an ionspray voltage of 5000 V. The capillary temperature was 300 °C. The MS recordings were carried out in selected reaction monitoring (SRM) mode with specific ion transitions of precursor ion to product ion at m/z 647.4→191.2 with collision energy (CE) of 38 eV, declustering potential (DP) of 85 V and entrance potential (EP) of 15 V for TY501, and at m/z 473.3→143.3 with CE of 23 eV, DP of 55 V and EP of 10 V for IS. The total analytical time was 5 min. PK analysis was performed using Phoenix Winnonlin. Initial estimates, AUC, and Cmax was determined using naïve pooling NCA. A one compartment model was fit to the data and PK parameters Ka, Vd, and CL were obtained from the models. Simulations were executed using the obtained parameters. All statistical significance was at the 95% confidence level.

### IR-MALDESI Imaging of the spatial distribution of vismodegib-word count 266

For MALDESI imaging, 2 replicate *G-Smo* mice were injected IP with 100mg/kg of POx- vismo or conventional vismodegib. Mouse brains were dissected free and placed on a foil barrier over dry ice for rapid freezing. 10 µm frozen sections of brains in the sagittal place were prepared in a cryotome, briefly thaw-mounted on glass microscope slides that had been uniformly coated with prednisolone as an internal standard using a pneumatic sprayer (TM-Sprayer, HTX Technologies, Carrboro, NC, USA), and then maintained at −10 °C on the sample stage of the IR-MALDESI source chamber prior to analysis. The stage translated the sample step-wise across the focused beam of an IR laser (λ = 2.94 µm, IR-Opolette 2371; Opotek, Carlsbad, CA, USA), which desorbed sample material from adjacent 100µm diameter sampling locations. An electrospray (50/50 mixture of methanol/water (v/v) with 0.2% formic acid) ionized the desorbed neutral molecules, and resulting ions were sampled into a high resolving power Thermo Fisher Scientific Q Exactive Plus (Bremen, Germany) mass spectrometer for synchronized analysis. The mass spectrometer was operated in positive ion mode from *m/z* 200 to 800, with resolving power of 140,000_FWHM_ at *m/z* 200. With high mass measurement accuracy (MMA) within 5 ppm maintained using protonated and sodiated adducts of diisooctyl phthalate as two internal lock masses at m/z 391.28428 and 413.26623, vismodegib and prednisolone were identified as protonated molecular ions [M+H^+^]^+^ at *m/z* 421.01695 and *m/z* 361.20095, respectively. To generate images from mass spectrometry data, raw data from each voxel was converted to the mzXML format using MSConvert software (*37*). These mzXML files were interrogated using MSiReader, a free software developed for processing MSI data (*38*).

### Bovine serum albumin (BSA) binding analysis

Vismodegib binding to BSA was evaluated by measuring decrease fluorescence emission spectra of tryptophan residues of BSA. Increasing concentrations of POx-vismo or conventional vismodegib were incubated in fetal bovine serum diluted with PBS to final BSA concentration of 40 mg/ml for 15 min at 25 °C. Emission spectra were recorded from 300 to 440 nm after excitation at 295 nm using SpectraMax M5 spectrometer. Both excitation emission slits were set at 1 nm. Presented data represents average of triplicate samples. To evaluate the release of vismodegib and binding to BSA over time, POx vismo or conventional vismodegib samples were added to 250 µL of FBS and incubated at 37 °C for predetermined time points. After the incubation, the samples were transferred to a prewarmed 10 kDa MWCO Vivacon filter tube and spun at 12,000xg for 10 min at 37 °C. Supernatants were passed through Strata™-X 33 µm polymeric reversed phase column then filtrates were mixed with acetonitrile and analyzed on HPLC as described. Presented data represents average of triplicate sample.

### *In vivo* toxicity studies

Toxicity of POx-vismo and conventional vismodegib was evaluated in healthy wildtype C57BL/6 mice. POx-vismo was injected IP and conventional vismodegib was administered either IP or by oral gavage to groups of 3 replicate mice on days P10-P12 and then every over day until P21. As mice at P12-21 are expected to gain weight steadily, we used the weights of age matched littermate controls to define the expected weight at each time point. Mice consistently weighing less than 90% of the controls were considered to be growth impaired.

For bone toxicity studies, healthy WT mice at P10 were injected IP with 100 mg/kg of either POx-vismo or conventional vismodegib at P10, P11, P12 and P14. CT images were taken on P15 using SuperArgus CT system. The scanning parameters for CT imaging were 70 kV, 300 µA, 39 msec, and 360 projections. CT images were imported into PMDO 4.0 and orthogonal views were selected so that 3 planes were visible. Mice were then euthanized, and femur were isolated, fixed in 4% PFA solution and the length of femurs was measured.

### Measurement of pharmacodynamic response by pRB quantification

Groups of 4 replicate *G-Smo* mice were injected with 150 mg/kg POx-vismo intraperitoneally (IP) and conventional vismodegib was administered by either IP injection (50- 100 mg/kg) or oral gavage (100mg/kg) on P12. 24h post injection, tumors from these mice were dissected free and dissociated as previously described (*39*). Dissociated cells were then treated with the Fix & Perm ® Cell Fixation and Permeabilization Kit per manufacturer instructions (Thermo Fisher Scientific). Fixed cells were incubated in 1:50 dilution of anti-phospho-Rb/Alexa Flour® 647 conjugated (Cell Signaling Technology) and 1:100 dilution of FX Cycle(™) Violet Stain for 2 hours in the dark on ice. Flow cytometry was then performed on an LSR Fortessa (BD Biosciences). Technical controls included no stain, single-stained and fluorescence-minus-one samples.

### Tumor pathology studies

Groups of 4 replicate *G-Smo* mice were injected IP on P10-12 with POx-vismo (100mg/kg) or conventional vismodegib at either 50mg/kg or 100mg/kg. After 24 hours, mouse brains including tumors were harvested, fixed, embedded in paraffin and processed for IHC as previously described (*40*) using antibodies to PCNA (Cell Signaling Technology, Danvers, MA, USA). Similarly, mouse brains including tumors were harvested from mice in the survival study as they reached the humane end point or P35 and processed for IHC. Stained slides were digitally acquired using an Aperio ScanScope XT (Aperio, Vista, CA, USA).

### *In vivo* efficacy studies

*G-Smo* mice were randomized into the indicated treatment groups with a minimum of 8 replicate mice per group and treated with either POx-vismo 100 mg/kg IP, conventional vismodegib at 50mg/kg administered either IP or orally and conventional vismodegib at 50mg/kg administered IP and compared to untreated controls. The indicated treatments were administered daily on P10-P12 and then every over day until P35, unless mice first developed symptoms of tumor progression. Symptoms of tumor progression included hunched posture, paucity of movement, ataxia and weight loss. All mice with symptomatic tumors were euthanized. The EFS was defined as the survival time until the development of symptoms. Survival times for each group were compared by Log Rank analysis.

### Statistical analysis

Statistical analyses for pharmacokinetic profile of vismodegib were performed using the two-tailed student’s t-test (Graphpad Prism, version 5.1.). For the survival analyses, the Log Rank test was used.

## Acknowledgments

We thank the UNC CGBID Histology Core supported by P30 DK 034987, the UNC Tissue Pathology Laboratory Core supported by NCI CA016086 and UNC UCRF and Ms. Yuling Zhao and Dr. Hedi Liu for assistance with histopathology, and the Chapel Hill Analytical and Nanofabrication Laboratory, supported by the National Science Foundation Grant ECCS- 1542015, for help with electron microscopy.

## Author Contributions

Conceptualization: DH, AVK, TRG, MSP; Methodology: EPR, JRK, AT, JDR, WZ, AVK, TRG, MSP; Investigation: DH, TD, CL, AT; Formal analysis: EPR, JRK, WZ, AT, JDR, TRG, MSP; Funding acquisition: AVK, TRG, MSP; Supervision: TRG, MSP; Writing – original draft: DH; Writing – review & editing: TRG, MSP. TRG, MSP.

## Data and materials availability

All data associated with this study are available in the main text or the supplementary materials.

