## Supplemental Information for "Poly(2-oxazoline) nanoparticle delivery enhances the therapeutic potential of vismodegib for medulloblastoma by improving CNS pharmacokinetics and reducing systemic toxicity"

**Supplementary Figures**

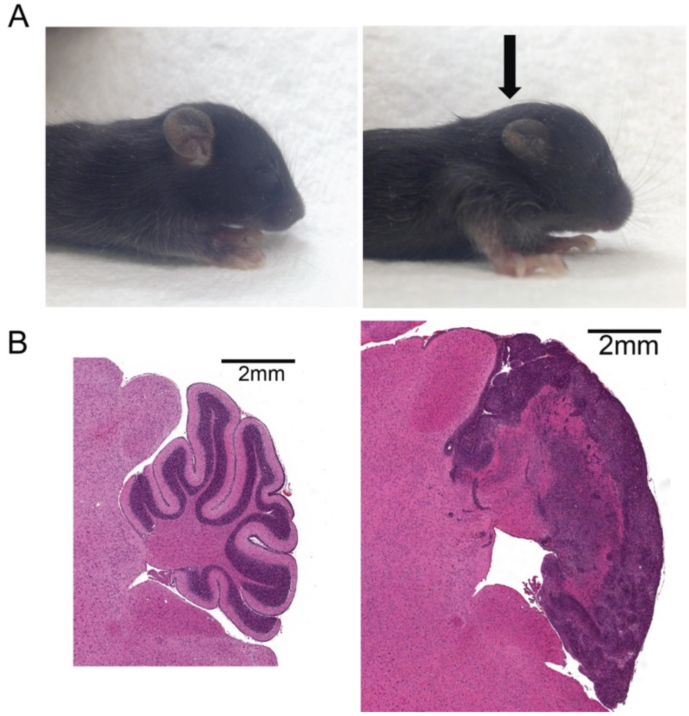


**Fig. S1.** Comparison of normal and *G-Smo* mice at P15. **(A)** Head shape and **(B)** H&E stained sagittal sections of the cerebellar region of normal (left) and *G-Smo* (right) mice.

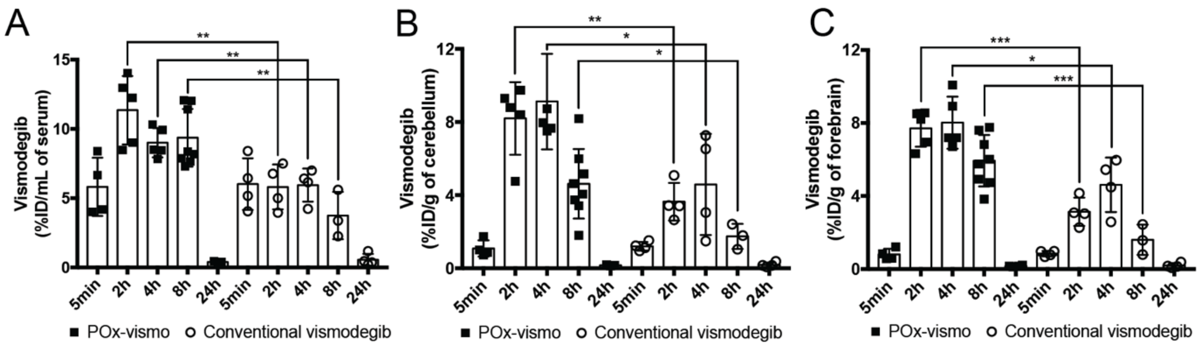


**D PK parameters**

| Tumor-bearing mice | | | | | | |
|---|---|---|---|---|---|---|
| Parameters | POx-vismo | | | conventional vismodegib | | |
| | Serum | Tumor | Forebrain | Serum | Tumor | Forebrain |
| AUC (0–∞) (ng*hr/mL) | 992869.75 | 574591.17 | 626488.79 | 706435.87 | 350346.09 | 351019 |
| Vd (mL) | 4.73 | - | - | 11.41 | - | - |
| CL (mL/hr) | 0.683 | - | - | 0.809 | - | - |
| **Wild-type mice** | | | | | | |
| Parameters | POx-vismo | | | conventional vismodegib | | |
| | Serum | Cerebellum | Forebrain | Serum | Cerebellum | Forebrain |
| AUC (0–∞) (ng*hr/mL) | 1062953.1 | 644605.5 | 704583.85 | 540380.3 | 288539.5 | 269031.6 |
| Vd (mL) | 3.77 | - | - | 10.29 | - | - |
| CL (mL/hr) | 0.649 | - | - | 1.22 | - | - |

AUC(0–∞), area under the curve from time 0–infinite; Vd, volume of distribution; CL, total body clearance.

**Fig. S2.** Pharmacokinetic profile of POx-vismo and conventional vismodegib. Vismodegib concentrations in **(A)** serum, **(B)** cerebellum and **(C)** forebrain, 24 h after following single IP injections of POx-vismo or conventional vismodegib at 100 mg/kg, *p < 0.05, **p < 0.005, ***p < 0.0005. **(D)** PK parameters at given formulations from tumor-bearing mice and wild-type mice. For **A-C**, dots indicate data from individual replicates and error bars are the SEM.
